# Accounting for programmed ribosomal frameshifting in the computation of codon

**DOI:** 10.1101/293340

**Authors:** Victor Garcia, Stefan Zoller, Maria Anisimova

## Abstract

Experimental evidence shows that synonymous mutations can have important consequences on genetic fitness. Many organisms display *codon usage bias* (CUB), where synonymous codons that are translated into the same amino acid appear with distinct frequency. CUB is thought to arise from selection for translational efficiency and accuracy, termed the *translational efficiency hypothesis* (TEH). Indeed, CUB indices correlate with protein expression levels, which is widely interpreted as evidence for translational selection. However, these tests neglect −1 programmed ribosomal frameshifting (−1 PRF), an important translational disruption effect found across all organisms of the tree of life. Genes that contain −1 PRF signals should cost more to express than genes without. Thus, CUB indices that do not consider −1 PRF may overestimate genes’ true adaptation to translational efficiency and accuracy constraints. Here, we first investigate whether −1 PRF signals do indeed carry such translational cost. We then propose two corrections for CUB indices for genes containing −1 PRF signals. We retest the TEH under these corrections. We find that the correlation between corrected CUB index and protein expression remains intact for most levels of uniform −1 PRF efficiencies, and tends to increase when these efficiencies decline with protein expression. We conclude that the TEH is strengthened and that −1 PRF events constitute a promising and useful tool to examine the relationships between CUB and selection for translation efficiency and accuracy.

## INTRODUCTION

Across an organism’s genome, among those codons that decode for the same amino acids (codon families) some are used preferentially over others [1] (reviews in [2, 3]). The variation of this codon *usage bias* (CUB) within organisms’ genomes is mainly explained by selection for translation efficiency by protein expression demands –the translation efficiency hypothesis (TEH) [2, 3].

The main support for the TEH stems from two separate lines of evidence that concern associations between CUB and tRNA abundances and CUB and protein expression, respectively [3]. In the first line, studies revealed that the most frequently used codons –so called *preferred codons* [2]– typically match the most abundant iso-accepting tRNA in both *Escherichia coli* [4, 5] and *Saccharomyces cerivisae* [4]. Similar conclusions were drawn for a number of additional species [6, 7], but using tRNA gene copy numbers as proxies for tRNA abundances (based on the assumption that they correlate, as is the case in *E. coli* and *yeast* [8, 9]). Further studies suggest that employing these more abundant tRNAs in translation provides efficiency [10] as well as accuracy gains (at least for *E. coli* [11, 12] and *Drosophila melanogaster* [13]). Furthermore, exchanging preferred codons in highly expressed genes by non-preferred had strong effects on gene expression [14, 15]. If valid, these two mechanisms together imply that the disproportionate frequent usage of such preferred codons in an mRNA sequence is thus indicative of high translational efficiency.

The second line of evidence concerns the existence of associations between CUB and protein expression [16–20]. High protein expression demands are assumed to generate a selective pressure for translation efficiency [2, 21, 22]. Given the mechanisms of the first line of evidence, these efficiency increases could be provided by synonymous mutations that alter codon usage frequencies towards preferred codons [2, 23]. Taken together with the first line of evidence, these associations suggest that protein expression is also associated with translation efficiency, completing the support of TEH.

The test of TEH by detection of CUB-protein expression association does therefore not rely on a direct comparison between translation efficiency and protein expression. Instead, CUB indices serve as a measure for adaptation due to translational selection [19]. CUB indices are computed from a given mRNA sequence. Thus, the test of the TEH by CUB-protein expression association hinges on the assumption that the CUB of an analyzed mRNA appropriately reflects its translational efficiency.

The phenomenon of *programmed ribosomal frameshifting* (PRF) [24–30] indicates that the validity of that assumption may not always be warranted. Programmed ribosomal frameshifting is a process by which ribosomal translation of an mRNA sequence is induced to stall at specific sites, termed *slippery sites*, which leads to a rearrangement of the ribosome on the mRNA sequence [27, 31]. Translation then proceeds in a new frame that is shifted relative to the original *open reading frame* (ORF) [32]. This mechanism emerges in all domains of the tree of life [29, 33–35]. Here, we focus on events that shift ribosomes one nucleotide back in the translational direction, termed −*1 PRF* [27, 36], further restricting ourselves to *yeast*.

The function of −1 PRF differs across organisms. In viruses, −1 PRF is predominantly employed to package more information into available sequence material. By opening a new reading frame, the −1 PRF mechanism allows for dual coding. This function is exemplified in the *gag-pol* gene overlap appearing in many retroviruses [24], in particular human immunodeficiency virus (HIV) [37]. In these viruses, the appropriate proportion of *gag* versus *pol* protein expression is regulated by a −1 PRF signal [37–39]. In eukaryotes, −1 PRF is predominantly (> 99%) used for gene expression regulation [36, 40]. About 10% of genes in yeast are hypothesized to contain functional slippery sites [41]. Ribosomes redirected to −1 shifted frames majorly encounter *premature termination codons* (PTC), that is, stop codons that appear well before the poly-A tails of mRNA sequences [36]. This typically triggers the activation of the nonsense-mediated decay (NMD) pathway [42]. The NMD pathway induces the degradation of both the mRNA, as well as of the partly assembled protein, while leaving the ribosome intact [36]. Thus, −1 PRF acts as an NMD-mediated destabilizing element of mRNA [43–45]. −1 PRF efficiency appears to be controlled by sequence specific elements, such as miRNA [44], but can be uniformly affected across all PRF-harboring genes by mutations as well as drugs [45].

Thus, the reason why CUB may not appropriately reflect translation efficiency for genes containing −1 PRF signals, is because of concealed costs to translation. When a −1 PRF signal redirects a fraction of the mRNA-translating ribosomes to premature termination [29, 36], a translation efficiency cost is incurred that is not reflected in the sequence’s CUB index.

In this study, we retest the CUB-protein expression association that lends major support for the TEH by accounting for the −1 PRF phenomenon. We draw from *PRFDB*, a database of predicted −1 PRF signals in the *Saccharomyce cerevisiae* genome [41]. First, we devise a series of hypotheses for how −1 PRF signals are influenced by evolutionary pressures. In accordance with our hypotheses, we find evidence for the existence of a cost of −1 PRF mechanism maintenance. Second, we devise a general correction any codon usage bias index for mRNA containing −1 PRF signals. We use sequences from PRFDB to compute the new, corrected codon bias indices, and compare them to uncorrected values. We find that the TEH is robust against these corrections and strengthened under biologically plausible assumptions of a −1 PRF dependency on protein expression. We conclude that the −1 PRF signals offer untapped potential to analyze translation efficiency in mRNA.

## RESULTS

### The cost of PRF-1 maintenance

Effective −1 PRF events classically comprise three constitutive elements. The first element is a heptameric sequence –the *slippery site*-[24, 40]. This site has the structure (X XXY YYZ), where X can be any base (A,C,G,U), Y is either A or U, and lastly, Z is either A, C or U [31, 40, 46]. The second element is a *spacer sequence*, separating the slippery site from the third element [29]. The spacer sequence typically comprises around 1-12 nucleotides [30, 46]. The third element is a *pseudoknot*, an mRNA secondary structure that is thought to generate a mechanical tension on the spacer sequence when translating the slippery site [29, 30, 46]. To release the tension, the ribosome, while stalling at the slippery site, is pushed one nucleotide back [27, 29, 46]. The frequency with which the ribosome is redirected to the −1 frame is called −*1 PRF efficiency* [36, 45].

These criteria were used to generate an extensive database of −1 PRF signals, *PRFDB*, of the yeast genome by algorithmic search (see *Materials and Methods*). If PRFDB comprises a sufficiently large set of functional slippery sites, the effects of putative −1 PRF induced costs to translation efficiency should be detectable.

To this end, we devised three hypotheses. We derive these hypotheses from the assumption that the maintenance of −1 PRF signals carries an intrinsic cost to the organism. Furthermore, we assume that mutations that alter CUB are accumulated more slowly than mutations that alter −1 PRF efficiency, *p*. This is because the −1 PRF regulation mechanism is very likely to be affected by only a few mutations, whereas several mutations would be needed to significantly modify CUB. Hence, most short-term changes in protein expression demand in genes with slippery sites are more likely absorbed by −1 PRF.

We first introduce a framework to formulate these hypotheses (see Figure 1). Let L be the length (in nucleotides) of an mRNA sequence *x*. A single slippery site is located at site *l* of *x*, downstream from the start. Ribosomes are redirected to the −1 frame with probability *p*, the −*1 PRF efficiency*. They continue with the mRNA’s translation until encountering a premature termination codon (PTC) at a distance λ post-slippery site.

In a first hypothesis, we formed expectations about how the number of slippery sites per gene should vary with the ratio of protein expression to mRNA levels per cell. This ratio has been used as a measure for translation efficiency in past studies [10]. To formulate the first hypothesis, we conduct a thought experiment. Let us assume that an organism’s environmental conditions favor lower expression of a protein from a −*1 PRF*-gene. Then, increasing the average *p* in that gene will adapt the organism to the new conditions. Conversely, if such environmental changes demanded higher protein expression, genes containing −1 PRF signals may adapt by reducing *p*. If protein demands exceed the production capacity of a gene with very small *p*, the −1 PRF mechanism may offer only costs, but no benefits from protein regulation. Then, mutations that remove the heptameric slippery site signature will be selected for. Taken together, these effects imply an asymmetry in the probability of slippery site loss depending on the direction of protein demand changes. Losses of slippery sites are likelier when protein demands increase than when they decrease. −1 PRF signals should thus become rarer with increasing protein expression levels.

**FIG. 1.**
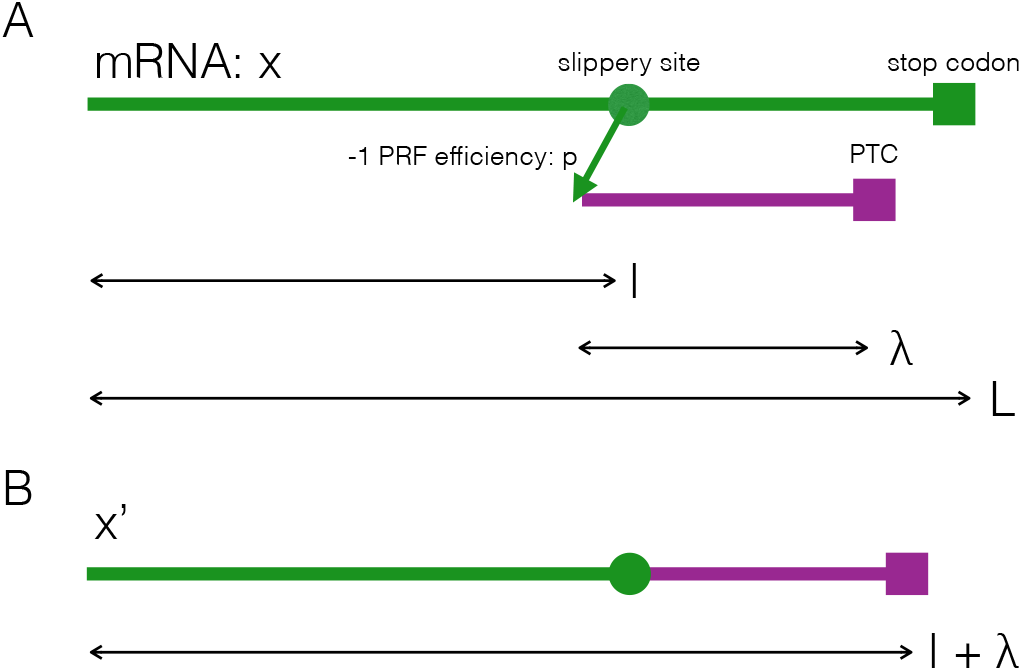
Notation for −1 PRF signals. A) The mRNA sequence in which a −1 PRF signal does not lead to ribosomal frameshifting is termed *x* (green). Arrows indicate the distances in nucleotides. B) The mRNA sequence with −1 PRF is termed *x*′. This is the concatenation of the mRNA *x* until the slippery site (green), and the −1 frame after the slippery site until a premature termination codon is encountered (violet).

The second hypothesis states that the costs of −1 PRF maintenance are reduced as slippery sites approach the 5’ end or start of the mRNA. Every time ribosomes slip through −1 PRF, the energy of translating a stretch of length *l* + λ of mRNA is wasted for protein production. Since λ is roughly constant across within-gene sites (see Fig. S1), the cost to translational efficiency incurred from −1 PRF should depend on *l*. Therefore, diminshing *l* will minimize cost. Thus, the distribution of slippery site positions relative to the gene length should be skewed towards mRNA starts, that is, small *l* values.

This effect is expected to be weaker than the effect in the first hypothesis. In the first hypothesis, if protein demands exceed yields generated by a gene and *p* is small, a fitness cost is incurred from i) the reduction of protein expression, ii) the cost of NMD-mediated mRNA degradation and iii) the cost of unnecessary translation of frameshifted mRNA sequences. Displacing the slippery site would only reduce the latter cost. Thus, we expect the signal for this hypothesis to be weaker than for the first hypothesis.

The third hypothesis states that a skew in the distribution of the location of the slippery site within a gene (relative to that gene’s length) should also become more pronounced with higher expression levels. This is because for fixed p, increasing expression levels should mirror an increase in −1 PRF induced translation costs. Thus, the benefits of slippery sites closer to translation-initiation positions should increase with expression levels. As in the second hypothesis, selection for such an effect is likely to be very weak compared to effects in the first hypothesis.

Figure 2 shows the results of testing all three hypotheses. Figure 2A shows that −1 PRF signals in genes become rarer with gene expression, consistent with our first hypothesis. Figure 2B shows that slippery sites positions are more prevalent in the first half of a gene than in the second, consistent with our second hypothesis. The frequency of sippery site diminishes towards both extremes of mRNA sequences, but more markedly so towards the stop codon of the mRNA. This skew persists when including all genes from the PRFDB (see Figure S2). Figure 2C shows that the average slippery site position is displaced towards the 5’ end of the mRNA as protein expression levels increase from 10^2^ molecules per cell to 10^5^ molecules per cell. Above 10^5^ molecules per cell, the uncertainty around the averages becomes large due to low sample sizes, and unambiguous deductions become impossible. The decreasing trend in the average *l*/*L* with protein expression is confirmed by a linear regression. Lastly, we retested all of the hypotheses with an alternative integrated data set from PaxDB, and obtained the same results (see Figure S3).

Thus, with increasing demand for protein production, the data suggest that production gains will primarily be attained by disposing of the PRF-1 mechanism present in a gene, rather than minimizing the cost of faulty translation. We speculate that, most likely, this occurs by altering the slippery site sequence. The test of the other two hypotheses involving slippery site displacement (or loss of large-*l* slippery sites) provide further evidence for an intrinsic −1 PRF cost to translation. The selective pressure on this latter adaptive process appears to be weak.

### Retesting the translational efficiency hypothesis while accounting for −1 PRF genes

Since −1 PRF signals carry a translational cost, their presence in an mRNA sequence changes the meaning of the associated CUB index. In fact, a slippery site may indicate a mismatch between the CUB index of that sequence and the actual efficiency with which it is translated (see Fig. 3).

To elucidate this point, consider an mRNA without −1 PRF signal. In the −1 PRF signal’s absence, the CUB index of a translated mRNA sequence appropriately reflects how efficiently ribosomes elongate it in a pre-specified time (see Fig. 3A) [10, 11, 13]. However, given a functional slippery site in that same mRNA sequence, the interpretation changes (see Fig. 3B). Let us assume an average −1 PRF efficiency *p*. Then, a fraction *p* of the times in which a ribosome is in the process of translating a specific mRNA, it will be redirected to another frame by −1 PRF. This redirection leads to degradation of the mRNA. Thus, the production of the same number of proteins will require more energy and time in the presence of a −1 PRF signal than it would in its absence. However, traditional CUB indices will give no indication of that process. If accounted for, −1 PRF must therefore lead to a downward correction of the mRNA’s translational efficiency and with it, of the associated CUB index.

**FIG. 2.**
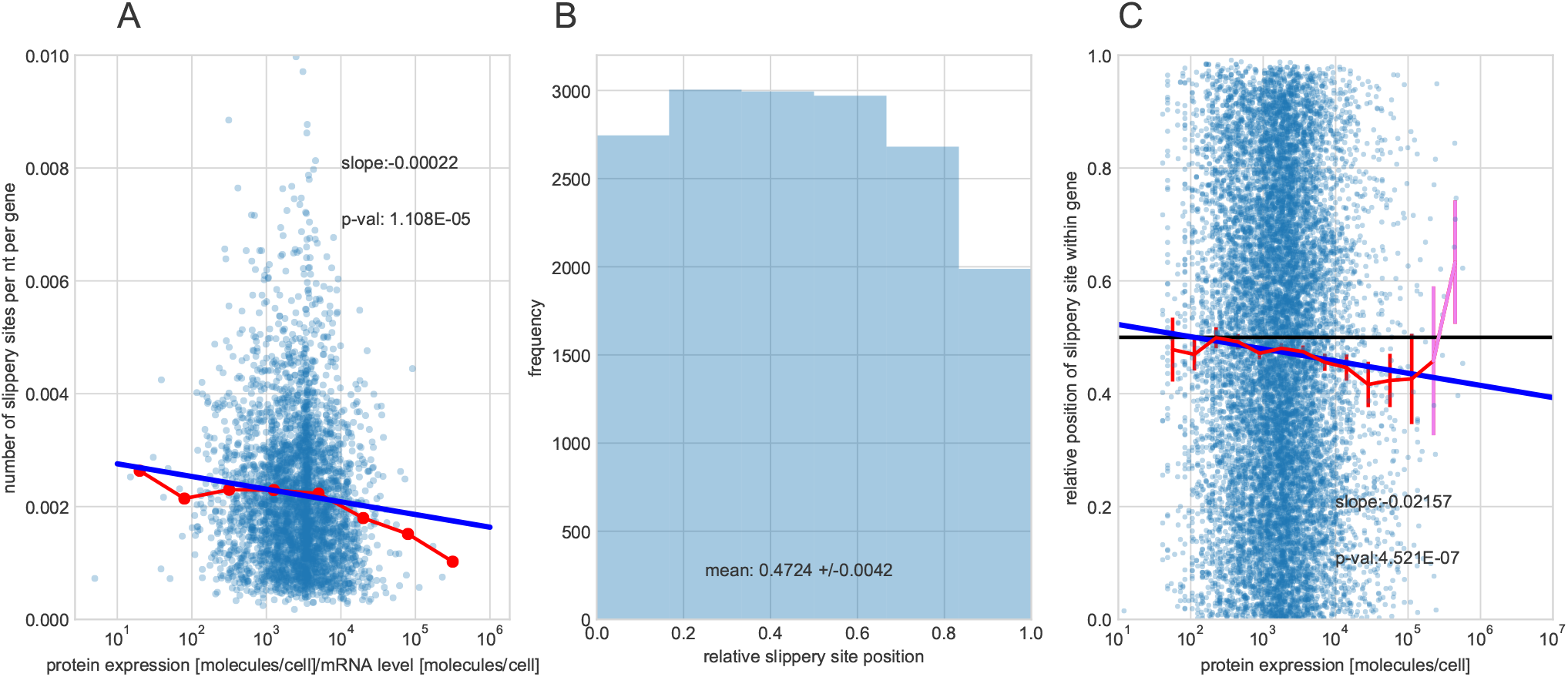
Testing the cost of −1 PRF mechanism maintenance. A) Number of slippery sites —identified by relying on *Nupack* algorithm for pseudoknot detection [47]— per gene and per nucleotide across protein expression levels per mRNA level for genes from the von der Haar data set [48]. The red dotted line is the average number of slippery sites per gene. The blue line is a regression line through the data set. The text in the panels gives the slope of the line and the p-value of the t-test for a non-zero slope value. B) The frequency distribution of the within-gene positions of the slippery sites, relative to the length of the gene, *l*/*L*. To ensure comparability, only genes from the von der Haar data set are considered. The mean of the distribution, as well as its 95% confidence intervals, are smaller than 0.5 -the expectation in the absence of selective pressure. C) Slippery site positions relative to gene length across protein expression levels in the von der Haar data set. The red line is the average slippery site position (computed across 20 bins of equal width in logarithmic scale) and the widgets are the uncertainty (±1.96 standard error of the mean) around the average estimate. Averages with large uncertainties are in violet. Analogously to A), a regression line with the corresponding slope and p-value are added.

Since a substantial fraction of genes may contain −1 PRF signals (hypothesized to be ≈ 10% of genes in yeast [41]), neglecting this effect might introduce considerable biases in CUB index values. These biases could affect the associations found between CUB and protein expression [16-20]. To address this issue, in the following we propose two general corrections for CUB indices in the presence of −1 PRF signals, and retest the basis for the TEH stemming from CUB and protein expression correlations.

### Codon Usage Bias Index Correction from Translation Efficiency

In a first approach, we derive an estimator for a CUB index correction from basic relationships between CUB indices and translation efficiencies of an mRNA. In the absence of a −1 PRF signal, the CUB on *x* will be measured by some index function *I*(*x*). As noted, if a functional −1 PRF signal is present, *I*(*x*) does not account for the translation efficiency loss due to the unproductive translation of the hybrid sequence *x*′ (see Fig. 1). We aim to derive a ‘‘corrected” index *I_c_*(*x*) that more appropriately reflects the diminished translation efficiency in the presence of −1 PRF signals.

To find an expression for *I_c_*(*x*), we begin with the assumption that there exists a monotonous mapping *F* that maps the translation efficiency *η*(*x*) of a sequence *x* (in the absence of −1 PRF) to a codon usage bias index *I*: *I*(*x*) = *F*(*η*(*x*)). *F* is a monotonously increasing function, where increases in translation efficiency are reflected by increases in the codon usage bias index. Research by Tuller et al. further suggests that *F* is concave [10], such that codon usage bias index values saturate with increasing translation efficiency values. We define translation efficiency *η*(*x*) classically as an input-output energy ratio. More specifically, *η*(*x*) is the ratio of the per-protein energy *E_p_*(*x*) contained in the *n*(*x*) synthesized proteins from *x* to the energy, *E_i_*(*x*), exerted into producing those proteins in a sufficiently long time frame Δ*t*:

**FIG. 3:**
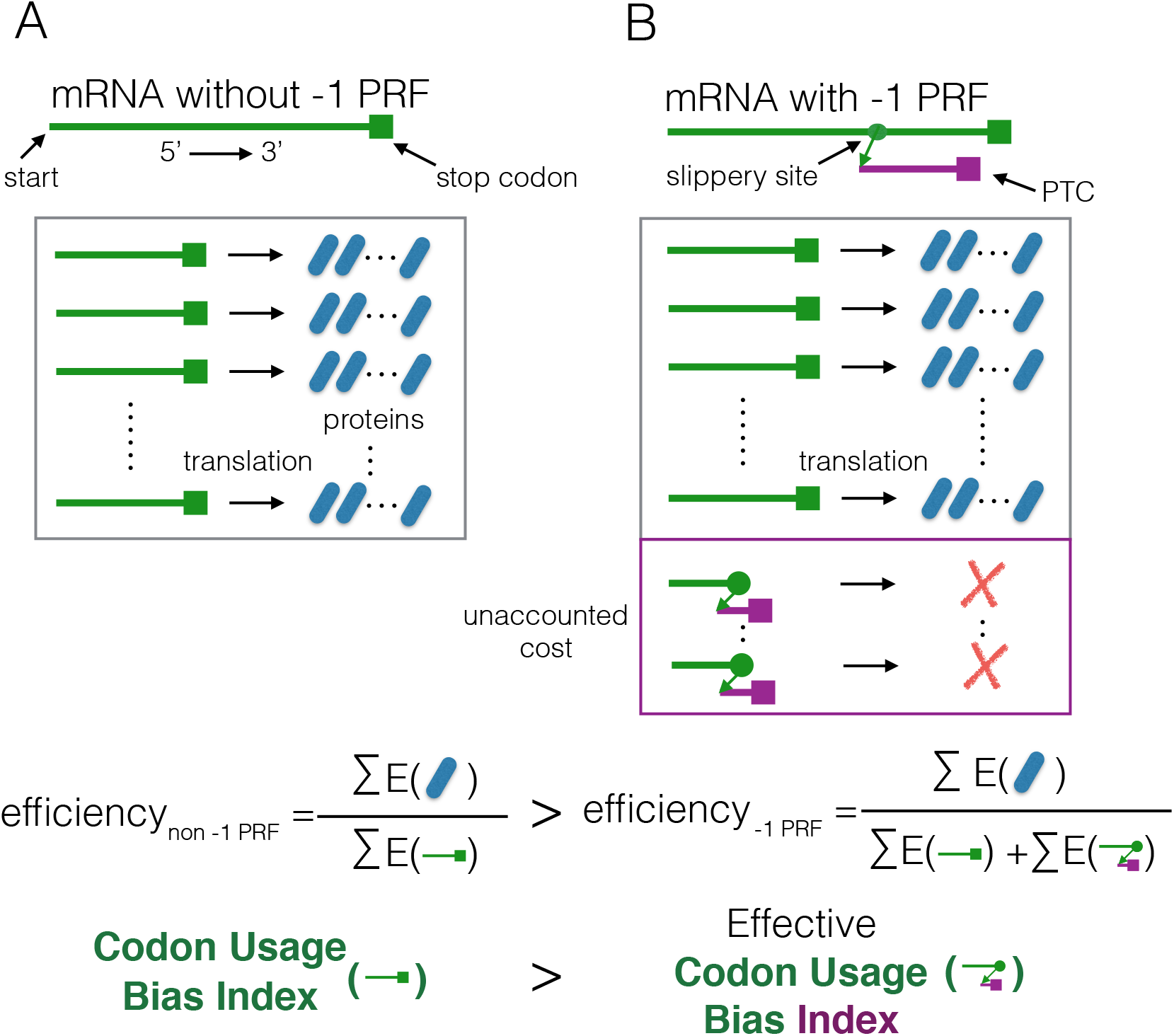
-1 PRF signals bias codon usage bias indices’ predictiveness of translation efficiency. A) An mRNA sequence (green line) without a −1 PRF signal. Within a given period of time, the transcribed mRNA in the cell will produce a finite number of proteins. We define the efficiency of the protein synthetisation process is defined as the energy content of the proteins (the nominator) divided the mRNA transcription and translation processes (the denominator) [10]. The value of a codon usage bias index reflects the efficiency of the protein synthesis process. B) The same mRNA considering a slippery site (green point). The slippery site redirects ribosomes to a −1 shifted frame, where they encounter premature termination codons (PTC). The same number of proteins are produced as in A). An additional cost, unaccounted for in A), is incurred for protein production due to NMD-mediated protein decay. Thus, the effective translation efficiency when accounting for −1 PRF must be smaller than A). This diminished efficiency should be reflected by a correction in the codon usage bias index value.

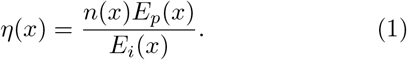

Here, the energy going into the synthetization machinery, *E_i_*(*x*), does not contain the expenditure for −1 PRF’s regulatory use. Thus, for a sequence *x* that additionally carries a functional slippery site, we have

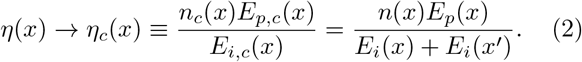

Here, the same number of proteins were produced as when −1 PRF was not considered, that is, *n_c_*(*x*) = *n*(*x*).

Since the final proteins are structurally equivalent to those in the absence of −1 PRF, we also have *E_p,c_*(*x*) ≡ *E_p_*(*x*). However, more energy was expended to produce these proteins, and therefore *E_i,c_*(*x*) = *E_i_*(*x*) + *E_i_*(*x*′) > *E_i_*(*x*). Their synthetisation requires at least *E_i_*(*x*). A part of the expended energy on translation is not implemented in the proteins, but in the translation of *x*′ and the NMD pathway activation, *E_i_*(*x*′). Thus, it follows that *η_c_*(*x*) < *η*(*x*).

The corrected usage bias statistic *I_c_*(*x*) is defined as

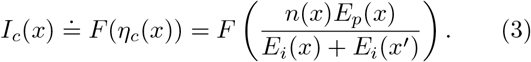

Since the form of *F* as well as the values of *E_p_*(*x*), *E_i_*(*x*) and *E_i_*(*x*′) are either difficult to measure or unknown, we aim to to compute *I_c_*(*x*) indirectly from *I*(*x*). To this end, we separate the translation efficiency *η_c_*(*x*) into two components:

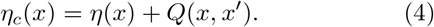

Solving for *Q*(*x,x*’) gives

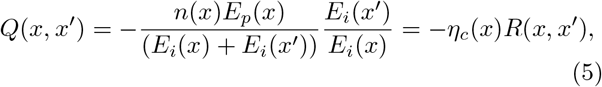

where 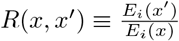. With this, we have

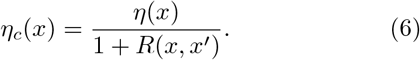

Further analysis of the ratio *R*(*x, x*’) is complicated by the inability to directly measure *E_i_*(*x*) and *E_i_*(*x*′). To address this issue, we pursue an approach where the ratio is approximated by information about the relative one-elongation energies spent translating *x* and *x*′. More specifically, we assume that each time *x* is translated, an energy input of *T*(*x*) ≐ *E_i_*(*x*)/*n*(*x*) is expended per time unit Δ*t*. The total number of times translation is initiated on the mRNA sequence, *N*(*x*), is *N*(*x*) = *n*(*x*)/(1 — *p*). This is because *n*(*x*) corresponds to the fraction 1 – *p* of times that an mRNA-elongating ribosome remains in frame. Analogously, the number of times translation is interrupted by a −1 PRF event is 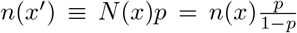. Hence, each time the sequence *x*′ is translated, an energy expenditure of *T*(*x*′) ≐ *E_i_*(*x*′)/*n*(*x*′) is ensued. Note that *n*(*x*′) does not correspond to a protein number. The ratio between these two is approximated by:

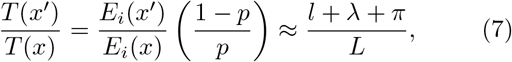

which is independent of the number of synthesized proteins *n*(*x*). Here, we have assumed that ratio of the per-translation attempt energies expended for the non-frameshifting to the frameshifting scenarios correspond roughly to the lengths of the translated sequences *x*′ and *x*, respectively. However, there is an extra cost to each frameshifted mRNA, *x*′ stemming from NMD-mediated degradation. This cost is accounted for by *π*, which is an unknown fixed cost associated with each −1 PRF event. *π* is measured in units of equivalents of translation cost per nucleotide.

With (7), we can approximate the *R*(*x,x*’):

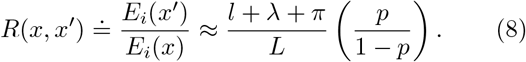

With this, we are ready to address the last approximation required to find an analytical expression for *I_c_*(*x*).

Since *F* is concave, it follows that *F*(*αη*) ⩾ *αF*(*η*) for 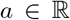. Thus, setting *a* = (1 + *R*(*x,x*’))^−1^, and using definition (2), we find a lower bound for *I_c_*(*x*):

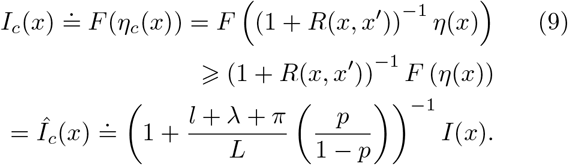

The approximation for *R*(*x, x*’) results in reasonable and useful properties of *η_c_*(*x*). More specifically, as *p* → 1 we have that *η_c_*(*x*) → 0. This entails that no mRNA is translated into proteins, as expected. The same follows if *π* → ∞. The extraction of *a* from within the brackets leads to the undesirable behavior that as *p* → 1 or *π* → ∞, we have that *Î_c_*(*x*) → 0. Instead, *Î_c_*(*x*) should approximate some minimal value *I*_*c*,min_(*x*) = *F*(0). A major drawback of *Î*_*c*_(*x*) is its reliance on *p* and *π*, which are both unknown. Typically, *p* is assumed to lie between 1 – 10%, but may reach up to 70% (for example the *EST2* gene in yeast, [45]). It is unclear how large *π* should be. In the following, and if not stated otherwise, we assume that *π* = 0 to give a conservative estimate of the effects of-1 PRF on the TEH.

#### Codon Usage Bias Index Correction from Averaging

In a second approach, we define a correction for a −1 PRF-aware CUB index using a balancing principle. To this end, we add the CUB index *I*(*x*) in successful, non-NMD mediated translations of *x* and the index *I*(*x*′) of untranslated *x*′, while weighing both with their respective probability of occurrence:

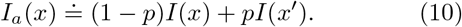

The corrected index *I_a_*(*x*) is thus the expected value of *I* when considering that *x* is translated a fraction (1 – *p*) of the time, and *x*′ is translated a fraction p of the time. *I*(*x*′) therefore acts as a penalization function. *I_a_* may underestimate the extra energy costs to −1 PRF incurred from the activation of the NMD-mediated degradation processes, since these are not comprised in *I*(*x*′). Similarly to *Î_c_*(*x*), the major drawback of the estimator *I_a_*(*x*) is its dependence on *p*. As before, we thus evaluate *I_a_*(*x*) for different, plausible values of *p* to assess −1 PRF’s effect on the TEH. However, *I_a_*(*x*) has the advantage that it is also well defined for *p* = 1.

#### Reexamining the TEH with corrected CUB indices

Both corrections *Î_c_* and *I_a_* represent a change in the value of a codon usage bias index given a −1 PRF index.

The models for correcting *I* presented here do not relate these measures to protein expression levels, *P*. They do therefore not predict protein expression. Instead, they allow us to reexamine the association between protein expression and a corrected codon usage bias index that in part underpins the TEH.

To examine how these corrections affect associations between CUB and protein expression, we used the widely employed *codon adaptation index* (CAI) [49] as an example for a CUB index *I*. We then computed both corrected *I_c_* and *I_α_* for different values of *p* and for all genes in the von der Haar data set [48] that contain −1 PRF signals. For *I_c_* we also assumed different values of *π*.

A precise computation of *I_c_* and *I_a_* would require information about how *p* varies with protein expression levels. Since no such associations have been identified as of now, we follow two approaches. First, we treat all −1 PRF sites equally, assuming an equal *p* value for all, independent of the expression levels *P* of the genes they are located in. Second, we allow *p* to slowly decrease with the *P* of the gene.

Additionally, we have found no evidence that the value of *π* is dependent on any characteristic of mRNA, such as its length, or codon composition [29, 42]. NMD is an evolutionarily conserved surveillance pathway [42]. Its activation may thus differ in the energy expenditure between organisms (indeed, in *yeast* NMD does not require an exon-junction complex, unlike most other eukaryotes). However, we have found no indication that the energy required to fully complete the NMD process will substantially vary from mRNA to mRNA within an organism. Thus, we assume that *π* is uniform across genes, although we do not know how large it is relative to other energy inputs.

Figure 4 shows that correcting for the presence of −1 PRF signals with a −1 PRF efficiency of up to *p* = 0.3 does not substantially affect the CUB index to protein expression relationship for both CUB index corrections, if the extra cost of mRNA degradation by NMD is neglected (*π* = 0). Figure 4A) shows that all CAI values for all genes are diminished when using the correction *I_c_*. A substantial correlation between protein expression and corrected CAI index remains. Accounting for the frameshifted sequences *x*′ also diminishes codon adaptation measures *I_a_* considerably compared to uncorrected CUB index values, as shown in Figure 4B). However, unlike with *I_c_* this reduction does not correspond to a uniform negative offset. Instead, the effect of accounting for −1 PRF is to both shift and broaden the distribution of *I_a_*(*x*) values relative to *I*(*x*) (see Figure S4).

The two corrections *Î_c_* and *I_a_* show different sensitivities to the −1 PRF efficiency. In particular, the sensitivity of *Î_c_* to −1 PRF efficiency is mediated by the value of *π*. Figure 5A shows that the correlations between both *I_c_* and *I_a_* to protein expression levels decline for increasing values of *p* and different values of *π*. As *p* increases, and for low costs of NMD-mediated mRNA decay, the correlation of *I_c_* to protein expression levels declines to very low values (~ 0.2 at *p* = 0.9 for *π* = 0,10,100). If the energetic cost of −1 PRF induced mRNA degradation, *π*, becomes larger (*π* = 1000, 10000), the correlation declines very rapidly with *p*, and almost vanishes when *p* reaches ~ 0.5. For *I_a_* estimates, correlations’ dependency on *p* is moderate, never falling below 0.4 at biologically implausible values of *p* of unity.

Since the data in Figure 2 strongly suggest a cost to −1 PRF maintenance, we also explored how a −1 PRF efficiency decline with protein expression levels *P* consistent with such cost would affect the correlations in Fig. 5A. Unlike a uniform *p* across protein levels, corrections to *Î_c_* are expected to be larger at low expression levels than at high levels. The −1 PRF decline is assumed as follows: *p*(*P*) = *p_b_*/log_10_(*P*), where we call *p_b_* the *baseline −1 PRF efficiency*. Figure 5B shows that under this assumption, correlations between protein expression and *Î_c_* rises with larger baseline −1 PRF efficiencies for all except the largest values of *π*, while again *I_a_* remains unaffected. We observe analogous results using the Spearman rank correlation (see Figure S5).

These results corroborate the support for the translation efficiency hypothesis. For uniform −1 PRF efficiencies and for *Î_c_*-based estimates, correlations of corrected CAI to protein expression levels decline with *p*. They only become sufficiently diminished to challenge the TEH when both the values of *π* and *p* are very high. Indeed, according to our mathematical framework, *π* = 1000, 10000 corresponds to the cost of either translating a large gene, or tenfold that cost. Except for *π* = 10000, all correlation values between corrected CUB indices and protein expression levels remain at around 0. 4 — 0.5 for biologically relevant −1 PRF efficiencies of *p* =1 — 10%. For *I_a_* estimates, correlation seems generally robust to changes in *p*.

When −1 PRF efficiencies decline with protein expression -in accordance with a cost to −1 PRF-most correlations between *Î_c_*-based estimates and protein expression rise with −1 PRF baselines. Except for *π* = 10000, all correlation values increase with −1 PRF baselines. This adds support for the TEH, suggesting that −1 PRF conceals stronger associations than measured with uncorrected codon usage bias indices.

## DISCUSSION

In this study, we have investigated the hypothesis that −1 PRF presupposes translational costs to an organism, while at the same time generating benefits associated with protein expression regulation. We explored whether such a cost might be identified more directly in data and used both PRFDB [41] and the von der Haar data set [48] to address this question. We devised three hypotheses for likely signals of such cost in −1 PRF carrying genes. We could not find contradictory evidence for any of those hypotheses in the data. In a second step, we explored whether these costs, if not accounted for, bias impor tant measures of CUB. We devised two new general approaches to correct CUB indices for the presence of −1 PRF signals. We then tested whether the concealment of such costs may unduly influence, falsify or strengthen, one classical test of this translational efficiency hypothesis: the association of CUB indices with protein expression levels. Under the assumption of uniform −1 PRF efficiencies, the energetic costs related with NMD activation would need to be implausibly high to warrant this conclusion. We find that on the contrary, assuming that −1 PRF efficiency decreases with protein expression levels -as suggested by the existence of a cost to −1 PRF-, the TEH is strengthened. Thus, taken together, our results suggests that high organismal demands for specific proteins are reflected in CUB-mediated translation efficiency gains.

**FIG. 4.**
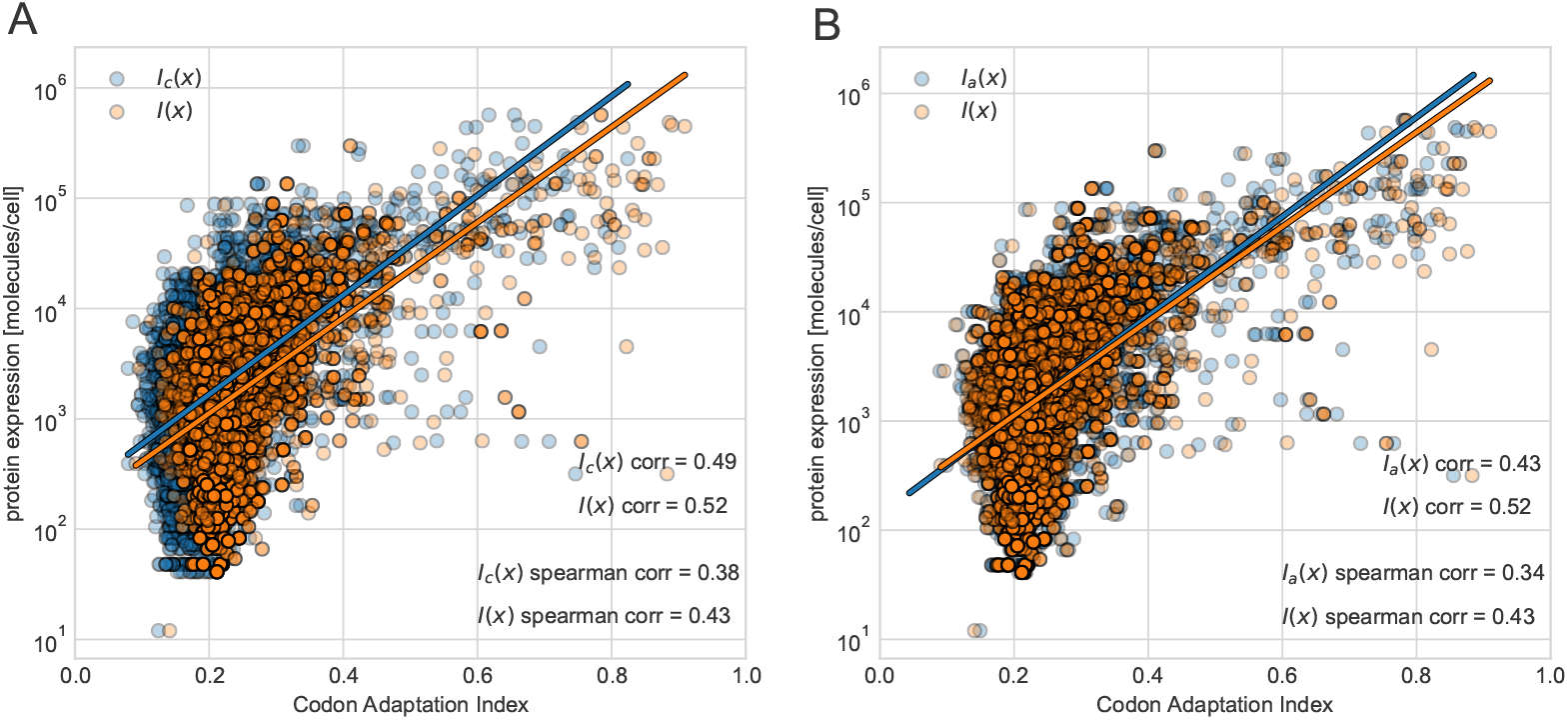
Corrected and original codon adaptation indices versus protein expression levels for genes containing algorithmically identified −1 PRF signals and with known expression levels [16], assuming *p* = 0.3 and *π* = 0. A) The original codon adaptation index values of mRNA sequences *x* of genes from the [16] data set are shown as orange, semitransparent points. The shown genes all contain algorithmically identified −1 PRF signals. The shape of the relation between CAI and protein expression is similar to that shown in [16] for all genes with known expression levels. Blue, semi-transparent points denote the values of −1 PRF corrected CAI values, *Î_c_*, versus same protein expression levels. Regressions between the log_10_-protein expression levels and both corrected an original CAI values are shown as orange and blue lines, respectively. B) is analogous to A) using the corrected CUB index *I_a_*.

**FIG. 4.**
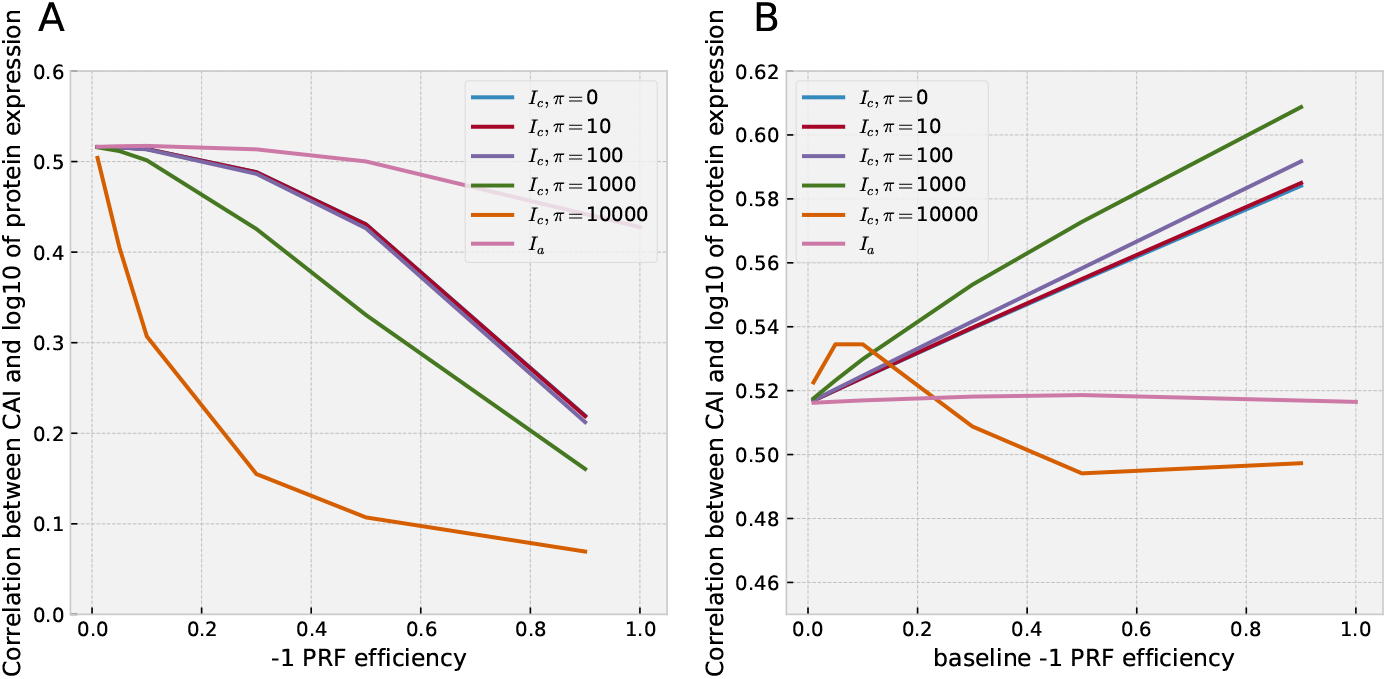
Correlation values between corrections of CAI to −1 PRF presence (*Î_c_,I_a_*) to protein expression levels *P* [48] for different −1 PRF efficiencies, *p*. A) For *Î_c_*, we considered different values of *π*, the cost of degrading an mRNA after −1 PRF, namely *π* = 0,10,100,1000,10000. For *Î_c_*, correlations values were computed at −1 PRF efficiencies *p* = 0.01,0.05,0.1,0.3,0.5,0.9. *Î_a_* was computed at the same levels except for *p* = 0.9, where instead *p* =1 was used. B) Analogous to A), but with *p* dependent on protein expression. For each gene, the −1 PRF efficiency is computed by *p*(*P*) = *p_b_*/log_10_(*P*), where for *p_b_*, the *baseline −1 PRF efficiency*, we chose the same values as for *p* in A).

Our study comes with a series of caveats. A first caveat of this study lies in that the slippery sites are inferred (utilizing the *Nupack* algorithm [47] for mRNA pseudoknot detection), but not always experimentally confirmed [45]. Previous studies suggest that inferred slippery sites are likely to be functional. In [45] it is reported that in previous work, 9 out of 9 high confidence −1 PRF sites detected by methods used in the PRFdB database in *S. cerivisae* were confirmed to be functional in vivo. Crucially, whether a −1 PRF site is regarded to be *functional* depends on whether it exceeds a predetermined p-threshold. For example, in [45], a search of −1 PRF slippery sites identified 10 candidate genes in *EST2*, three in *EST1*, 2 in *STN*, and 1 in *CDC* in mRNA involved in telomerase. Out of these, and employing a cutoff of 1% for −1 PRF efficiency, seven carry functional slippery sites (*EST2*:3, *EST1*: 2, *STN*:1, *CDC*:1) (see [45], Table 1). Had a cutoff of 0% −1 PRF efficiency been employed, all except one slippery sites would be functional. This limited sample provides confidence that the methods described by [41] appropriately capture biological mechanisms.

Even with such uncertainty, it is unlikely that the presence of non-functional −1 PRF sites in the analyzed data would affect the claims on an intrinsic −1 PRF maintenance cost (Fig. 2). These claims would only be biased if the probability of a candidate site to be functional were affected by either protein expression levels of the gene within which it resides or alternatively, the relative position in the gene of the putative slippery site. Note that these effects should arise from mechanisms that are independent of the ones studied here. Again, we are unaware of any such mechanisms acting in yeast, except for the requirement of a spacer sequence. A spacer sequence of a minimum length of say, 8 nt, will prohibit slippery sites to be located within 8 nt of the mRNA end. This restriction will slightly bias the *apriori* position of candidate slippery sites. However, as average mRNA by far exceed this length, such a restriction cannot explain the effects documented here.

Another caveat of our study was the lack of estimates for *p* for the whole data set. Due to this restriction, we could not explore how −1 PRF efficiency *p* relates to CAI or protein expression. As assumed in Figure 5B, we expect that −1 PRF maintenance costs should on average translate into an inverse relationship between *p* and protein expression levels, because higher protein expression demands by the organism should be countered with reductions of *p*, minimizing the loss of mRNA to NMD-induced degradation. However, it is also possible that there exist cases in which −1 PRF has important functions in highly expressed genes. Hence, although there are fewer slippery sites for highly expressed genes and they are costly in terms of translation efficiencies, these sites could have large −1 PRF efficiencies. We have not found evidence in the literature to support this notion, and it would concomitantly contradict the evidence here presented. Additional testing of the hypotheses on the costs −1 PRF maintenance would be greatly helped if such information were available.

While our analysis of the behavior of *Î_c_* gives us an indication how −1 PRF costs to translation efficiency could bias CUB indices, these insights rely on the assumptions made during *Î_c_*’s derivation. Importantly, *Î_c_* is a lower bound to *I_c_*, which means that downward corrections of *I_c_* are potentially exaggerated. Thus, the claim that corrected CUB indices will leave the basis for the TEH unaffected is resistant to such a bias. Another key assumption is that CUB indices of an mRNA sequence should increase monotonically with translation efficiency. In practice, this association is surely not perfect, due to multiple additional influences on CUB from unrelated biological processes. For these processes to systematically bias our results they should i) dominate over translational efficiency effects or ii) result in directional effects when combined. Despite these possible shortcomings, the assumption reflects key properties with which many CUB indices are designed, namely to mirror translational efficiency. For example, the CAI uses ribosomal mRNA as a reference to compute preferred codons because it is highly plausible that they are translationally efficient.

Moreover, our results suggest that utilizing expected values for corrected CUB index definition, such as in the case of *I_a_*, is suboptimal. Analysis of the distribution of *I_a_* levels with *p* = 1 of genes in the von der Haar data set show that corrected codon adaptation can increase. This behavior contradicts the rationale behind introducing such corrections in the first place, and suggests that, surprisingly, *I*(*x*′) > *I*(*x*) can occur.

To test the hypotheses about the TEH derived here, we have only utilized genes with identified slippery sites, and not all genes. Testing on all genes would be indicated if the TEH was challanged. However, if the correlation coefficient measured within a subset in which a correction has been applied is not substantially diminished relative to uncorrected values, the same correction will not affect the whole set either. Therefore testing it on the whole set is not necessary.

The results indicating intrinsic −1 PRF maintenance cost and TEH support are in mutual agreement. −1 PRF maintenance induces a reduction of the size of *l* with increasing protein expression levels (Fig. 2C). At fixed *p*, as assumed in our analysis, significant reductions in *l* would diminish the penalization to the original, un-corrected CUB. This could only be compensated by increases of *p* or *π* with protein expression levels, contrary to intuition and available evidence [45]. In fact, our analysis shows that a more plausible *p* dependency to protein expression leads to a strengthening of the basic CUB index to expression level association.

These results have to be interpreted in the context of current codon usage bias research. The mechanism of *translational selection* (TEH) remains the main explanation put forward for selection based origins of codon bias. This explanation presupposes that silent mutations affect fitness via translation processes. These fitness increases originate from translation efficiency and accuracy gains, that in turn are hypothesized to stem from translation initiation and elongation processes. In fact, mRNA elongation rates do indeed appear to correlate positively with preferred codon frequencies [50], although better evidence would be desirable. However, for initiation processes -which are assumed to contribute the bulk of these fitness increases–, the effects of CUB on initiation rate increase remain subject of debate [20]. In fact, Tuller and Zur have analyzed the effect of the structure of the 5’ end of an mRNA on translation initiation and elongation rates and found various regulatory signals that affect these rates in different ways [51]. Indeed, the induced folding at the 5’ end of the ORF appears to affect with translation efficiency.

Overall, the evidence from codon usage bias statistics and its associations to tRNA abundance and protein expression offers a compelling narrative for the TEH. However, current research efforts aiming to identify the exact mechanisms that give rise to these associations must account for conflating selective forces. Indeed, the mechanisms laid out in the TEH may not be the only way in which selection shapes codon usage frequencies. For example, besides the abundance of tRNAs, other factors have been discovered to crucially affect elongation rates and hence, to be possible targets selection [21, 52-54]. More precisely, specific synonymous changes can influence mRNA splicing, mRNA secondary structure, protein stability as well as protein folding [21, 55-59]. Synonymous changes may also alter the *secondary structure of mRNA* and thus affect the rate of translation -as established in vitro [51, 60, 61] and subsequently in vivo [62]. A comprehensive review on how protein expression is fine tuned by codon usage bias is given by Quax [20]. Further, in a very recent study, a synonymous difference between mammalian cytoskeletal *β*- and *γ*-actin proteins was found to affect co-translational processing, ubiquitination, and co-translational degradation, leading to differential stability properties of the corresponding protein products [63].

Within genomes, the biologial role of a gene influences that gene’s CUB in various additional ways. Some codons are more abundant in genes depending on their function, displaying distinct codon bias patterns. Supek has reviewed how gene function modifies codon preferences [64]. Selective pressure to maintain (or alter) gene function is superimposed to what is expected from translational efficiency and accuracy optimization. Genes where altered CUB patterns have been found are involved in diverse functions: amino acid starvation responses, cyclical protein expression, tissue specific expression, cellular differentiation, stress responses, and carcinogenesis. While genes can differ in function, there are also differences in function within a gene’s sequence that also affect local CUB. For example, Tuller and Zur have surveyed the multiple roles of the 5’ end of coding sequences in gene expression regulation [51]. They hypothesize that due to multitude of regulatory signals found in that region, selection pressures regarding codon utilization are likely different than in other regions. Unlike cross-gene function, such effects are local, which should not affect our analysis. How effects from gene function may affect our results, crucially depends on their frequency and direction. The literature reviewed here does not appear to warrant the assumption that all the function-dependent selective pressures will align to influence CUB in the same way, leading to systematic bias. The phenomenon of −1 PRF is thus only one of many ways in which CUB may be influenced.

The particular appeal of the −1 PRF phenomenon lies in its potential to elucidate many of these processes. Because slippery sites are precisely localized and −1 PRF efficiencies measurable, −1 PRF signals constitute natural experiments to translation efficiency and accuracy theories. This is because the separation of the mRNA by a slippery site should create differential translational costs across that mRNA. This translational cost gradient should be reflected in codon usage bias differences. −1 PRF based approaches to analyzing codon usage bias behavior, like the one presented in this study, may thus offer novel tools to better understand the means by which translation efficiency gains are realized in nature.

## MATERIALS AND METHODS

### Slippery site data

We obtained a dataset of the *Saccharomyces cerevisiae S288C* strain genes with predicted −1 PRF events from the Programmed Ribosomal Fraemshifting database (PRFDB) of the University of Maryland [41]. These putatitve −1 PRF signals were identified using first a filter for slippery site identification, and subsequent detection of mRNA pseudoknot by means of the *Nupack* algorithm [47]. For a slippery site to be found, the a confluence of signals in mRNA is required. First, a site must match the pattern (X XXY YYZ), where X is some base, Y is either A or U, and Z is A, C or U. Second, a spacer sequence of a minimum of 8 nucleotides in length needs to exist between the slippery site and the following pseudoknot. Third, a pseudoknot predicted by minimum free energy values of the mRNA secondary structure is expected downstream the slippery site. The dataset also included a full list of gene annotations, accession numbers, the relative position of the slippery sites within the gene in which they were found, as well as the gene mRNA. For each slippery site, we computed were premature stop codons first appear downstream. With this, we computed the frame shifted sequences.

### Protein expression and mRNA level data

We obtained protein expression as well as mRNA level data from the Supporting Information of [48]. Von der Haar has produced an extensive curated data set that merges data from various sources. For protein expression, the data include the seminal studies of Ghaemmaghami et al. [16], Newman et al. [65], and Lu et al. [66]. Additional data stems from 46 further studies, specified in [48]. To ensure comparability, only studies were included in which yeast was grown in rich medium. Transcriptome data for mRNA levels were obtained from [67-71]. These data include about 6000 genes. The construction of the curated data set is described in detail in [48].

### Indices for codon usage bias

Many indices have been proposed to assess the degree of codon usage bias present in a gien mRNA sequence. Here, we focus on the codon adaptation index (CAI) [49, 72]. In the following, we give the implemented definition of the CAI. Let *L* be the length of the mRNA sequence in codons, c is the index of the synonymous codons decoding the same amino acid *a*, and *o_ac_* is the observed count of the synonymous codon *c* of amino acid *a* in the sequence. *C_a_* is the index set of all codons within an amino acid *a*. *A* is the index set of all amino acids *a*. We define

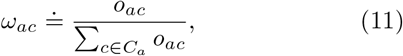

as the relative adaptiveness of codon αc within amino acid α of a given mRNA sequence.

Then, the codon adaptation index is defined as

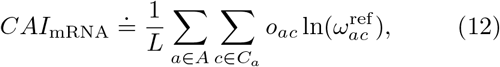

where *o_ac_* is measured on the observed on the mRNA of interest, and 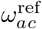 is taken from a reference sequence of highly expressed genes. For the reference sequence, we concatenated all of the mRNA of ribosomal genes [73]. The ribosomal genes were taken from the ribosomal gene database http://ribosome.med.miyazaki-u.ac.jp/ [74].

## ACKNOWLEDGMENTS

The authors thank Spencer Bliven and Lorenzo Gatti for stimulating discussions. This work was supported by the SystemsX – the Swiss Initiative for Systems Biology (grant number 51FSP0_163566).

## SUPPORTING INFORMATION FIGURES

**FIG. S1.**
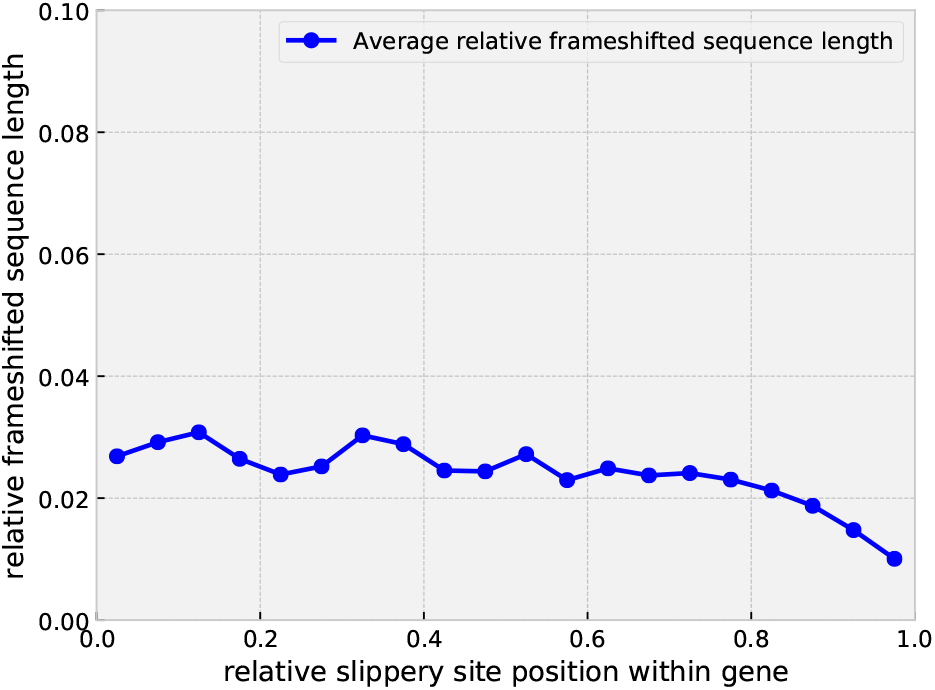
Average relative frameshifted sequence length, λ, across within relative gene length. Averages of the length of the frameshifted sequence after a slippery site are computed for 20 bins of equal width. Bins span the entire gene.

**FIG. S2.**
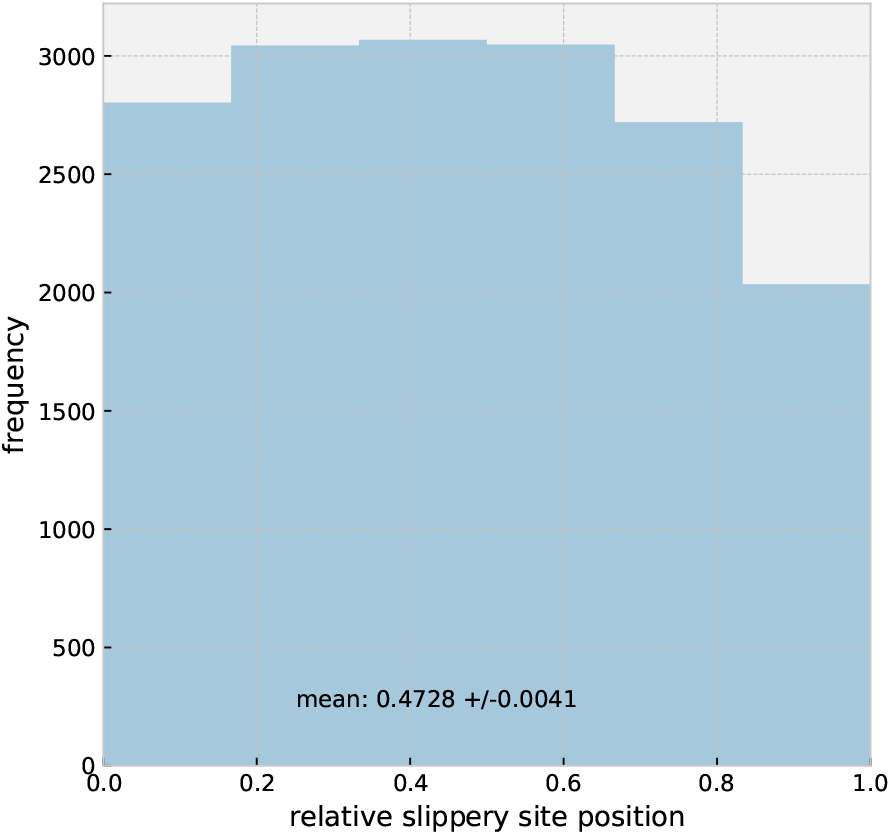
Frequency distribution of relative position of slippery sites *l*/*L* for all genes with −1 PRF signals in the PRFDB dataset. The mean of the distribution, as well as its 95% confidence intervals, are smaller than 0.5 -the expectation under randomly distributed slippery site locations.

**FIG. S3.**
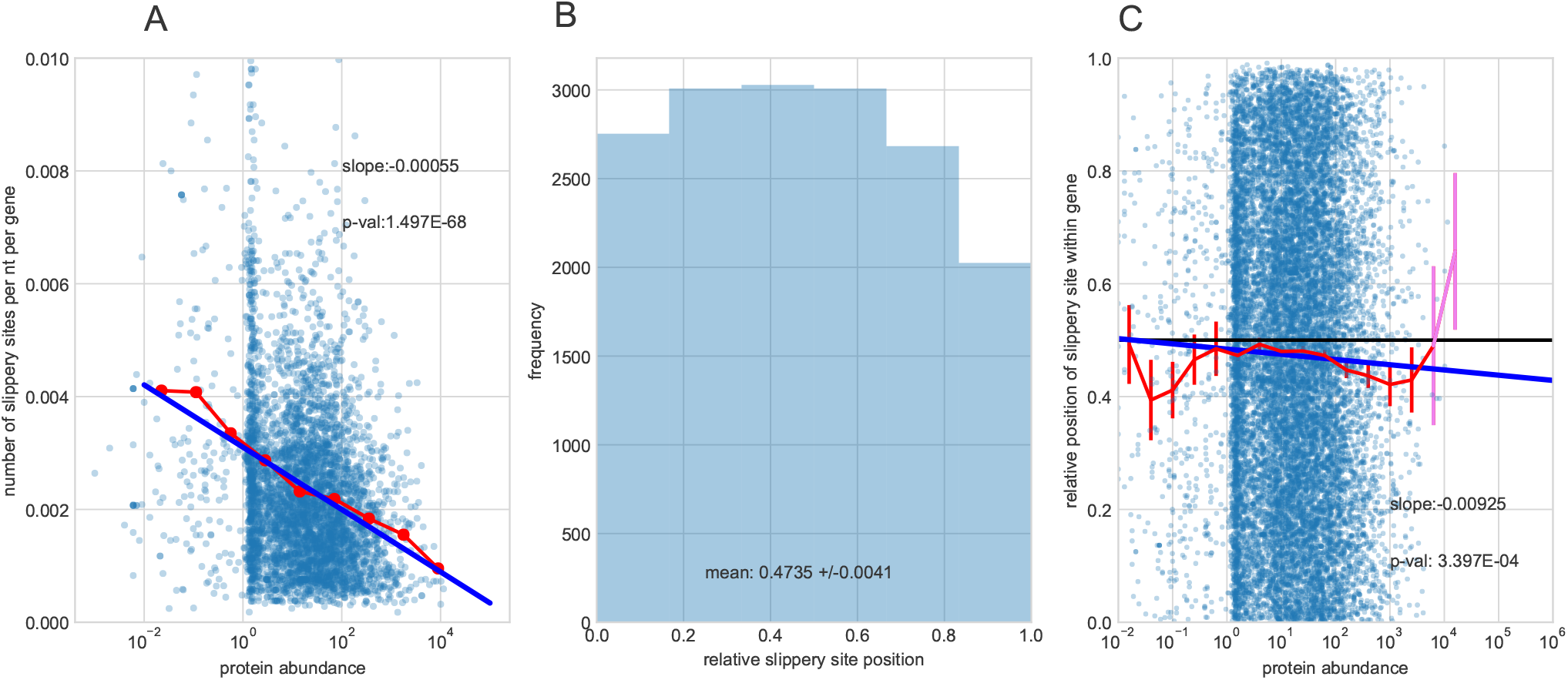
Testing the cost of −1 PRF mechanism maintenance. A) Number of slippery sites per gene and per nucleotide across protein expression levels (in molecules per cell) for genes from the integrated PaxDB data set. The red dotted line is the average number of slippery sites per gene. The blue line is a regression line through the data set. The text in the panels gives the slope of the line and the p-value of the t-test for a non-zero slope value. B) The frequency distribution of the within-gene positions of the slippery sites, relative to the length of the gene, *l*/*L*. To ensure comparability, only genes from the von der Haar data set are considered. The mean of the distribution, as well as its 95% confidence intervals, are smaller than 0.5 -the expectation in the absence of selective pressure. C) Slippery site positions relative to gene length across protein expression levels in the von der Haar data set. The red line is the average slippery site position (computed across 20 bins of equal width in logarithmic scale) and the widgets are the uncertainty (±1.96 standard error of the mean) around the average estimate. Averages with large uncertainties are in violet. Analogously to A), a regression line with the corresponding slope and p-value are added.

**FIG. S4.**
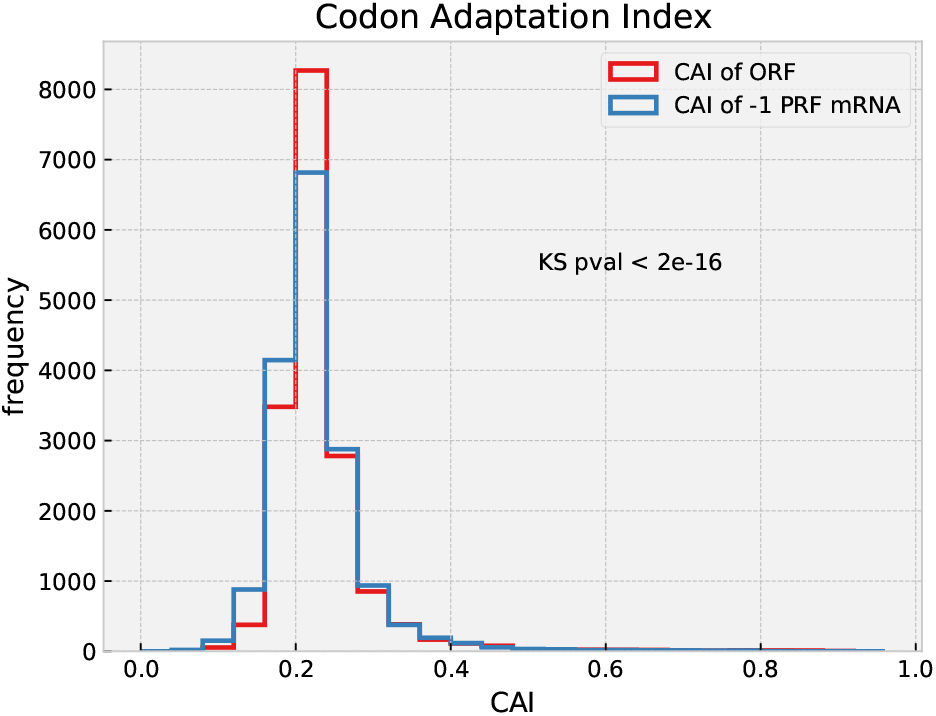
Codon Adaptation Index (CAI) frequency distributions of genes without −1 PRF signals (CAI of ORF) and with such signals (CAI of-1 PRF mRNA). The p-value given in the figure is for the two-sample Kolmogorov-Smirnov test for the two distributions.

**FIG. S5.**
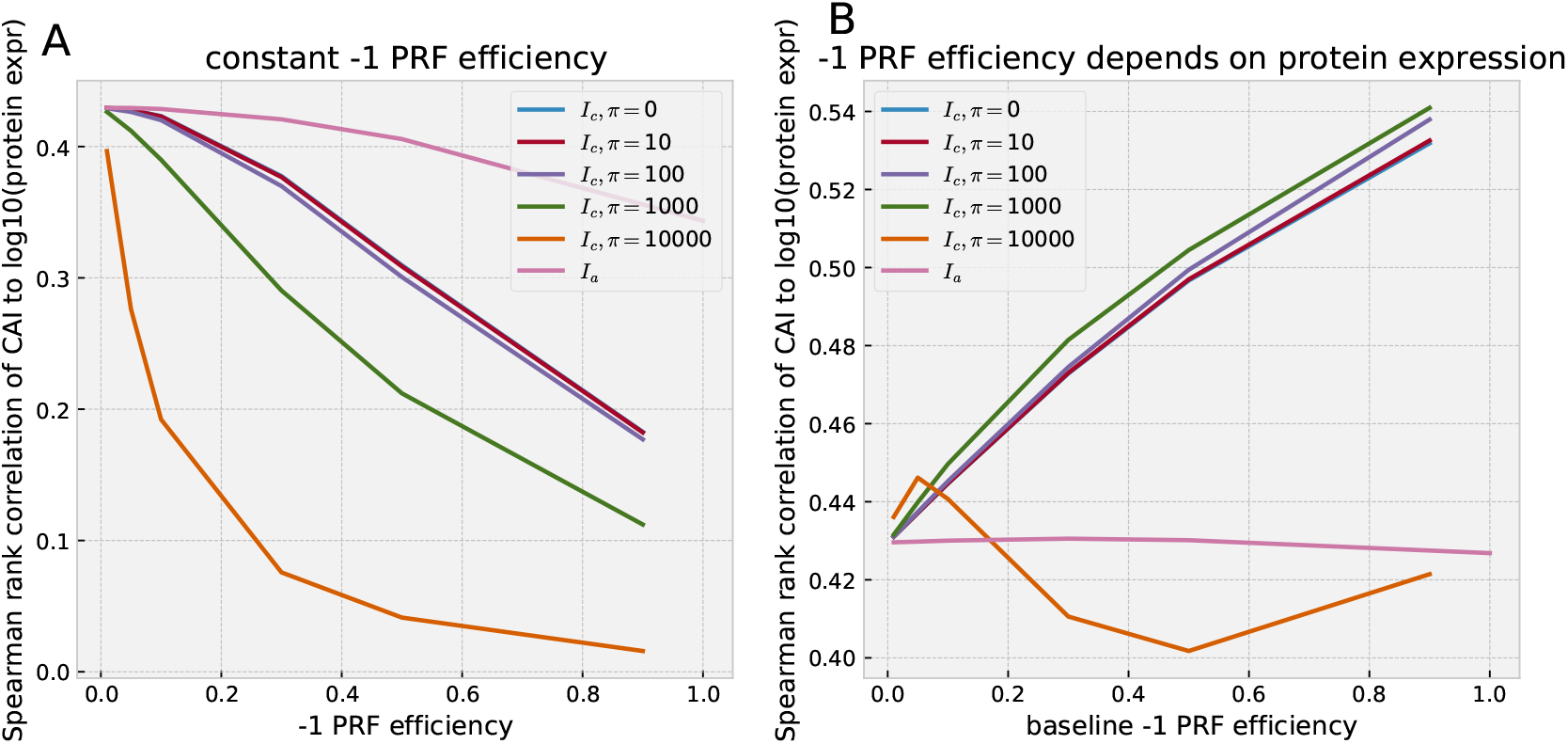
Spearman rank correlation values between corrections of CAI to −1 PRF presence (*Î_c_,I_a_*) to protein expression levels *P* [48] for different −1 PRF efficiencies, *p*. A) For *Î_c_*, we considered different values of *π*, the cost of degrading an mRNA after −1 PRF, namely *π* = 0,10,100,1000,10000. For *Î_c_*, correlations values were computed at −1 PRF efficiencies *p* = 0.01, 0.05, 0.1, 0.3, 0.5, 0.9. *I_a_* was computed at the same levels except for *p* = 0.9, where instead *p* =1 was used. B) Analogous to A), but with *p* dependent on protein expression. For each gene, the −1 PRF efficiency is computed by *p*(*P*) = *p_b_*/log_10_(*P*), where for *p_b_*, the *baseline −1 PRF efficiency*, we chose the same values as for *p* in A).

